# Fitness and transcriptomic analysis of pathogenic *Vibrio parahaemolyticus* in seawater at different shellfish harvesting temperatures

**DOI:** 10.1101/2023.05.03.539256

**Authors:** Zhuosheng Liu, Chao Liao, Luxin Wang

## Abstract

To better characterize the population dynamics of *Vibrio parahaemolyticus* (*Vp*) containing different virulence genes, two *Vp* strains were inoculated into seawater separately and incubated at temperatures (30 and 10 °C) mimicking summer and winter pre-harvest shellfish rearing seasons. The cellular responses of these two strains, one containing the thermostable direct hemolysin (*tdh*+) gene and the other one containing *tdh*-related hemolysin (*trh*+) gene, were studied at the transcriptomic level. Results showed that, at 30 °C, *tdh*+ and *trh*+ strains reached 6.77 ± 0.20 and 6.14 ± 0.07 Log CFU/ml respectively after 5 days. During this time, higher growth rate was observed in the *tdh*+ strain than the *trh*+ strain. When being kept at 10 °C, both *Vp* strains persisted at ca. 3.0 Log CFU/ml in seawater with no difference observed between them. Growth and persistence predictive models were then established based on the Baranyi equation. The goodness of fit scores ranged from 0.674 to 0.950. RNA sequencing results showed that downregulated central energy metabolism and weakened degradation of branched chain amino acid were observed only in *trh*+ strain not in *tdh*+ strain at 30 °C. This might be one reason for the lower growth rates of the *trh*+ strain at 30 °C. Histidine metabolism and biofilm formation pathways were significantly downregulated in both strains at 10 °C. No significant difference was observed for virulence-associated gene expression between 10 and 30 °C, regardless of the strains.

**SIGINIFICANCE:** Given the involvement of *Vp* in a wide range of seafood outbreaks, a systematical characterization of *Vp* fitness and transcriptomic changes at temperatures of critical importance for seafood production and storage is needed. In this study, predictive models describing the behavior of *Vp* strains containing different virulence factors are established. While no difference was observed at the lower temperature (10 C), *tdh*+ strain had faster growth rate than the *trh*+ strain. Transcriptomic analysis showed that significantly higher number of genes were upregulated at 30 °C than 10 °C. Majority of differentially expressed genes of *Vp* at 30 °C were annotated to functional categories supporting cellular growth. At the lower temperature, the down regulation of the biofilm formation pathway and histidine metabolism indicates that the current practice of storing seafood at lower temperatures not only protect the seafood quality but also ensure the seafood safety.

## Introduction

*Vibrio parahaemolyticus* (*Vp*) has been one leading seafood-borne pathogenic bacterium (gram-negative, rod-shaped) commonly found in marine environments, particularly in estuaries and coastal waters; it has been one major microbiological food safety concern for aquacultural products, especially raw oysters (1). To mitigate the risk of *Vp* contamination, shellfish farmers follow guidelines such as National Shellfish Sanitation Program (NSSP) in the United States for harvesting, handling, and storing oysters (2). Despite these control measures, over 36, 000 *Vp* infection cases associated with shellfish were annually reported in the United States, and recalls associated with *Vp* contaminated seafood products continue to occur (3–6). The undesirable public health consequences of *Vp*-contaminated food also bring economic burden. For instance, the total cost of illness associated with *Vp* infection increased from $40,682,312 in 2013 to $45,735,332 in 2018 in the United States (7).

Given its widely reported prevalence in aquaculture rearing environment and seafood products, efforts have been made to characterize behaviors of *Vp* in different stages from “sea to folk” (8, 9). Among different environmental parameters, temperature is one dominant factor significantly impacting the behavior of *Vp* (10). In coastal environment, significant lower levels of *Vp* or prevalence have been reported in winter months than summer (11, 12). This common finding was also accompanied by lowers *Vp* infection incidence rate in colder months based on the National Outbreak Reporting System (NORS) (13). These reported real-world evidence taken together underscored the correlation between ambient temperature and *Vp* prevalence and *Vp*-oriented Vibriosis.

RNA sequencing (RNA-seq) is a next-generation transcriptomic technique illuminating gene expression patterns in targeted organism, which can be applied to yield biological insights about physiological state of bacterial pathogens under different conditions (14, 15). Transcriptomic analysis has been used to investigate essential cellular mechanisms of *Vp* when surviving under simulated post-harvest practices (PHP) (e.g. cold storage, high salinity relaying, and acid-driven PHPs) (16–20). However, the physiological changes of *Vp* in natural seafood production environment, the impact of different virulence genes on its behavior, and how pre-harvest environment impacts the behavior of *Vp* during post-harvest handling and processing remain largely unknown. Therefore, this study aims to better understand the persisting mechanism of *Vp* in natural shellfish rearing environment, in particularly its cellular responses and virulence. This information, in turn, can support the development of novel control and monitoring strategies. The specific aims of this study were 1) investigating the survival and growth of *Vp* with different virulence genes in seawater at 30 and 10 °C and establishing predictive models for predicting *Vp* population under different conditions, and 2) profiling gene expression of *Vp* at 30 and 10 °C and identifying key changes in metabolic pathways and virulence factors.

## RESULTS AND DISCUSSION

### Primary models predicting *Vp* fitness in seawater at different harvesting temperatures

Populations of *Vp* stored in seawater at 10 °C or 30 °C over 10 or 5 days were enumerated based on the plate count method (**Figs. 1 and 2**). For the 10 °C trials, the inoculation level of *Vp* in seawater was 5.70 ± 0.06 and 5.76 ± 0.14 Log CFU/ml for *tdh*+ (ATCC 43996) and *trh*+ (ATCC 17802) strains, respectively. More rapid population decreasing was observed in the *trh*+ strain compared to the *tdh*+ strain. Continuous decreases of culturable *Vp* cells (ca. 2.0 Log CFU/ml) were observed from Day 0 to Day 7 and populational levels reached to 3.11 ± 0.19 and 3.57 ± 0.17 Log CFU/ml for *tdh*+ and *trh*+, respectively on Day 10. For the 30 °C storage trial, the initial inoculation level of *Vp* in seawater was 5.75 ± 0.07 and 5.74 ± 0.19 Log CFU/ml for *tdh*+ and *trh*+ strains, respectively. Growth of *tdh*+ and *trh*+ strains was observed and both strains reached the plateau phase with population levels at 7.11 ± 0.04 and 6.64 ± 0.08 Log CFU/ml respectively after 8 hours. After *Vp* reaching the plateau phase, significantly higher populational level in *tdh*+ compared with *trh*+ in persisted throughout the rest of incubation time. Difference in fitness between *Vp* strains containing different virulence genes has been reported by previous studies. S. Khouadja et al. (21) reported that *tdh*+ strain showed higher growth rate compared with *trh* strain when inoculated in sea bass serum and stored at 30 °C for 240 min.

**Figure 1.**
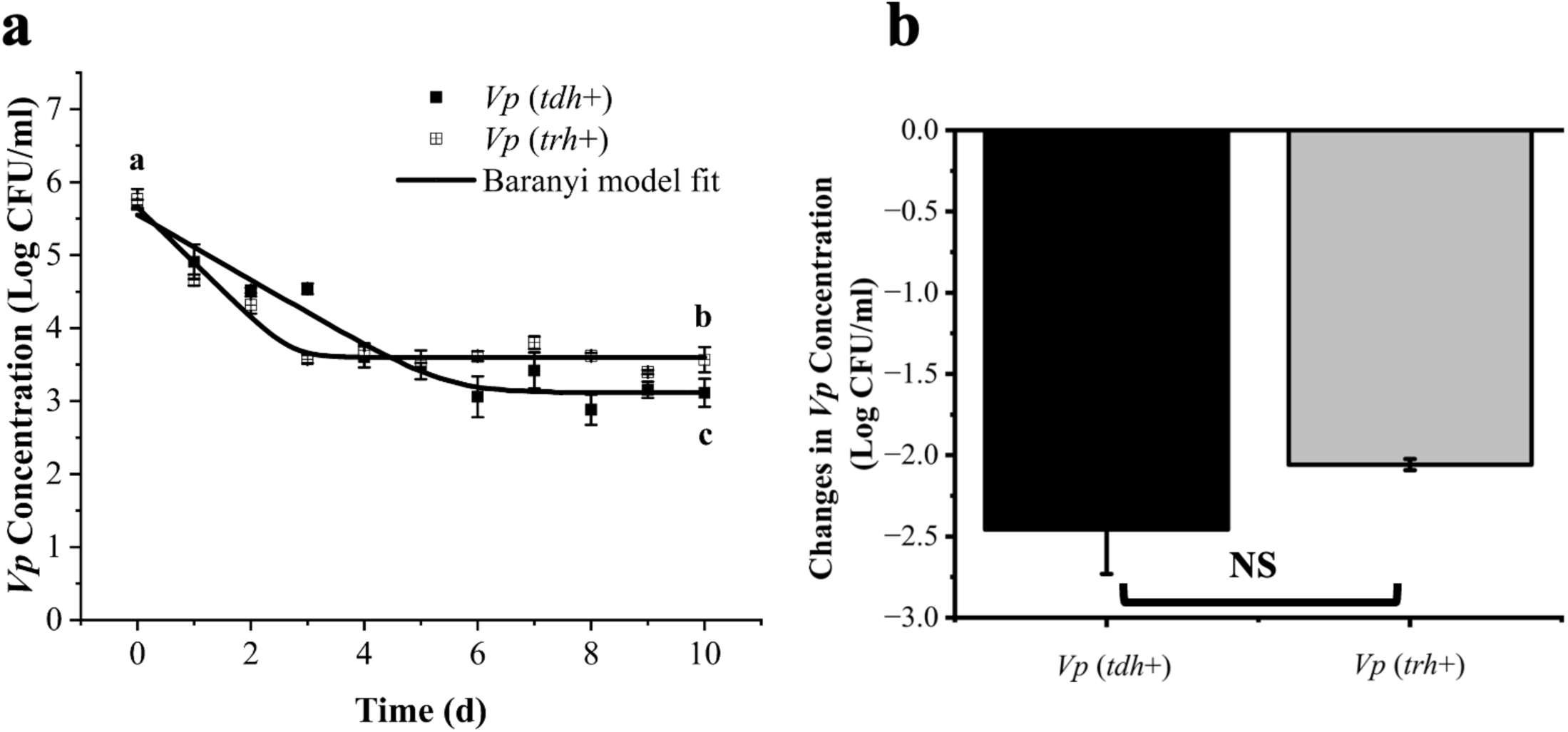
Primary predictive models of pathogenic *V. parahaemolyticus* strains (*tdh*+ and *trh*+) incubated in ater at 10 °C (a); and the comparison of the cell reductions of two strains on Day 10 (b). Different lower-case letters represent significant differences of bacterial counts between the *tdh*+ and the *trh*+ strains at different pling points. NS represents not significant.

**Figure 2.**
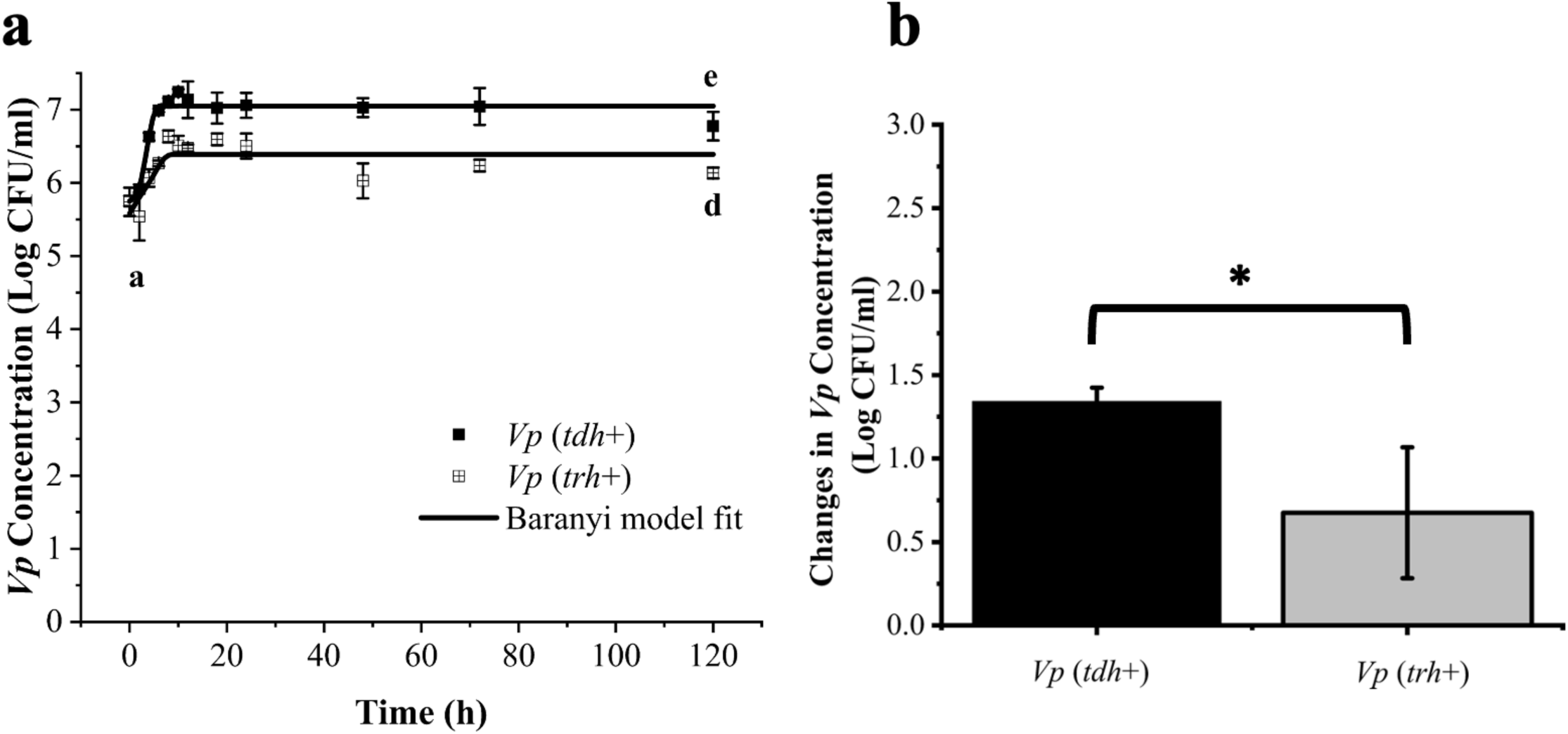
Primary predictive models of pathogenic *V. parahaemolyticus* strains (*tdh*+ and *trh*+) incubated in seaater at 30 °C (a); and the comparison of the cell reduction sof the two strains on Day 5 (b). Different lower-case letters represent significant differences of bacterial counts between the *tdh*+ and the *trh*+ strains at different sampling points. * represents a significant difference (*P* < 0.05).

The survival/growth data of *Vp* in seawater at 10 and 30 °C obtained from plate count results were further fitted by the Baranyi function to establish primary predictive models. The detailed parameters of Baranyi-based primary model predicting *Vp* fitness in seawater are listed in **Table 1**. The R^2^ of primary Baranyi models of *Vp* in seawater at 10 °C were 0.94 and 0.96 for *tdh*+ and *trh*+ strains respectively. At 10 °C, populations of the *Vp tdh*+ strain decreased from the initial values (IVs) to the final values (FVs) by 2.44 Log CFU/ml, meanwhile the *trh* strain decreased from the IVs to the FVs by 2.06 Log CFU/ml. The specific inactivation rate (SIR) in seawater at 10 °C was −0.45 ± 0.060 and −0.76 ± 0.11 Log CFU/day for *tdh*+ and *trh*+ strain respectively. No significant difference in *Vp* population reduction over 10 days at 10 °C was observed between *tdh*+ and *trh*+ strain based on difference between model predicted initial value and final value. C. Liao et al. (22) stored oysters inoculated with a five-strain *Vp* cocktail at 4.69 Log CFU/g at 10 °C for 11 days and reported SIR values of −0.073 ± 0.017 Log CFU/day. The difference in SIR values between the current study and the previous study might be caused by the different nutrient levels available in oysters vs. in seawater.

**Table 1.**
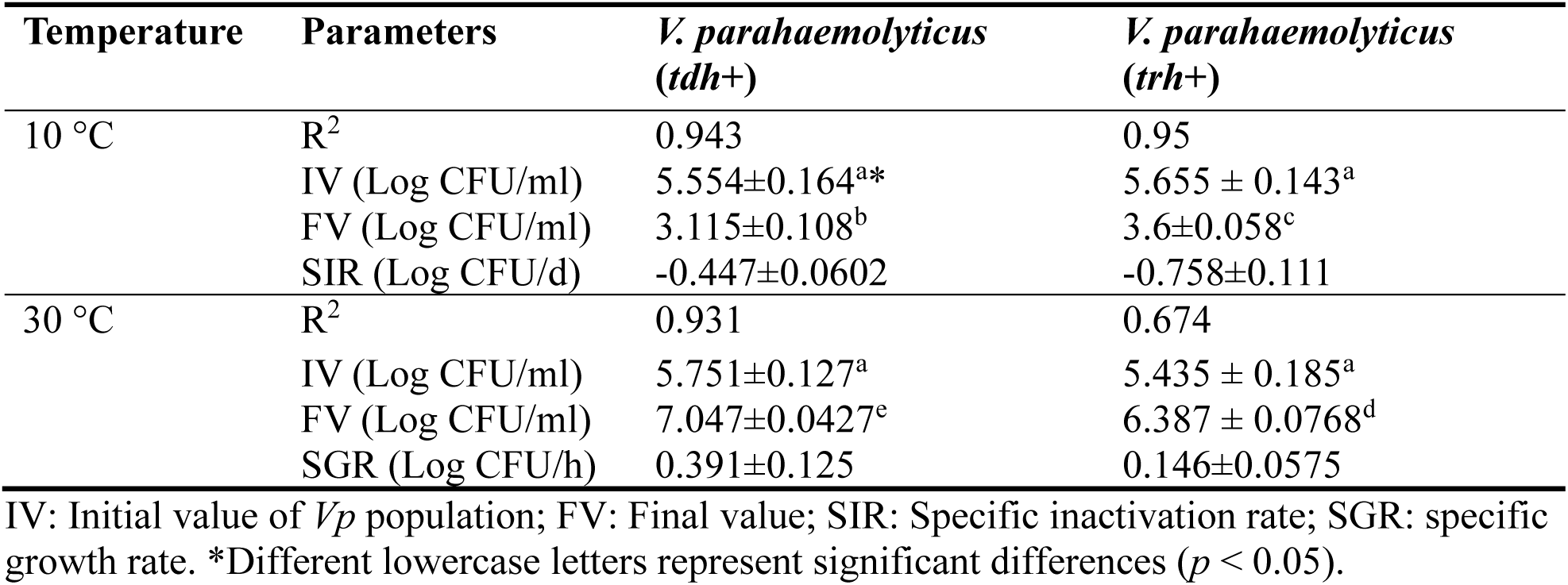
Primary predictive models predicting *Vp* fitness in seawater at 10 and 30 °C.

**Table 2.**
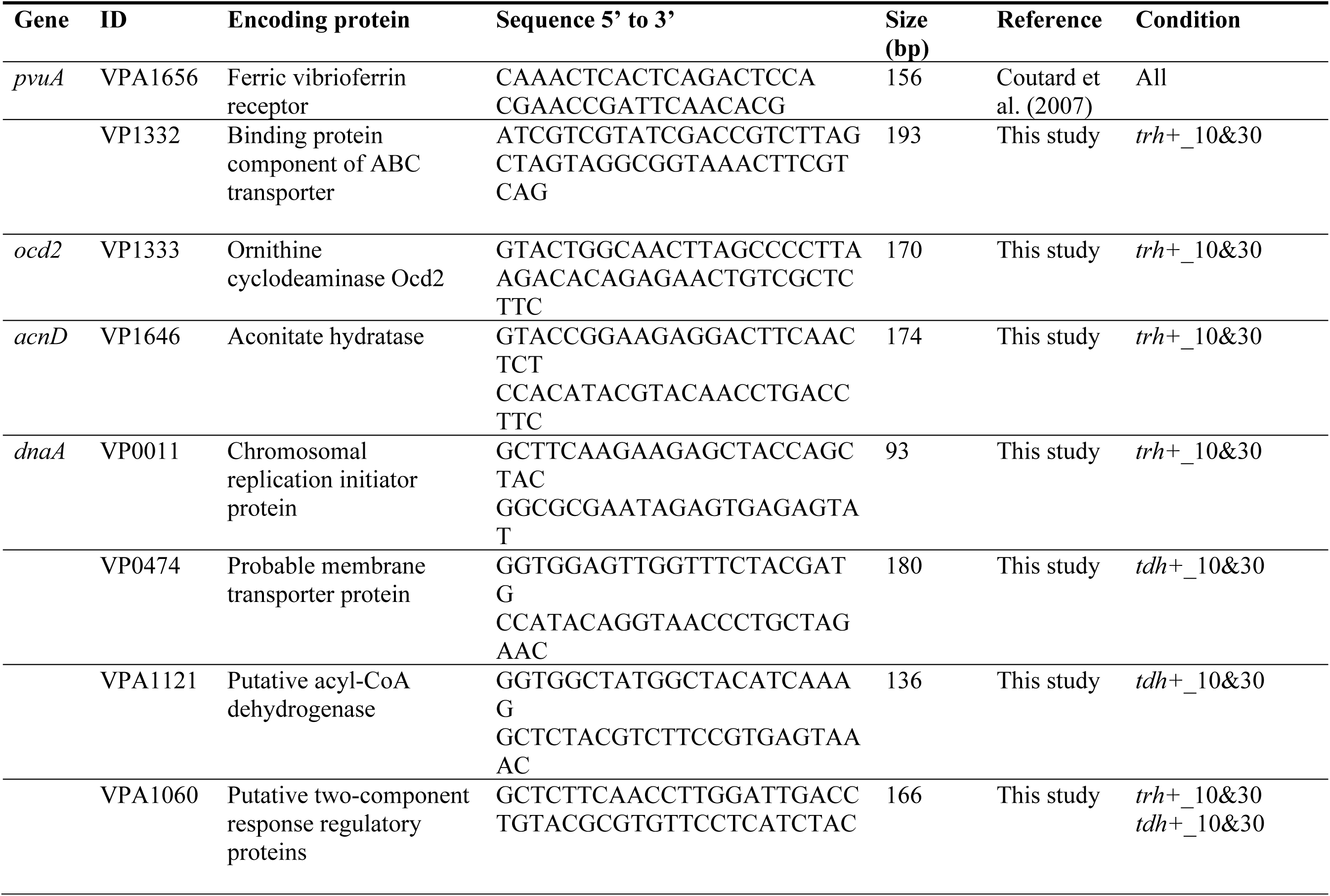
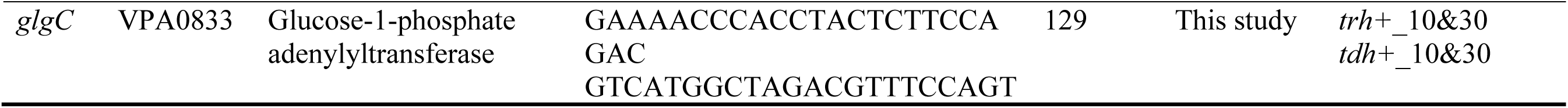
Primers used in qRT-PCR analysis for the validation of randomly selected differentially expressed genes identified by RNA-seq.

When the storage temperature was kept at 30 °C, the population of *tdh*+ strain increased from IVs to the FVs with a growth rate of 1.30 Log CFU/ml; the *trh+* strain increased from the IVs to the FVs with a growth rate of 0.96 Log CFU/ml. The R^2^ of primary Baranyi models of *Vp* in seawater at 30 °C were 0.97 and 0.95 for *tdh*+ and *trh+* strains, respectively. Higher specific growth rate (SGR) was observed on *tdh*+ (0.39 ± 0.13 Log CFU/day) than *trh*+ strain (0.15 ± 0.058 Log CFU/day) (*p* < 0.05).

### Transcriptomic profiles of *Vp* strains when surviving at different temperatures

Potential differences between the *tdh+* and the *trh+* strains at the transcriptomic level was further investigated at two temperatures. Raw RNA-seq reads were mapped to the reference transcriptome using Salmon. The mapping rate of aligning raw sequence reads with the reference transcriptome ranged from 65.30 to 78.30%. A total of 4,001 genes were successfully identified after the Salmon quasi-mapping against the protein coding sequence of *Vp* RIMD2210633. The Pearson correlation coefficient (PCC) of *Vp* gene expression profiles in the control (two hours after seawater inoculation) and the test groups (10 and 30 °C incubation over five days) was calculated to examine the linear relationship of gene expression patterns. As shown in **Fig. S2a,** PCC of *Vp* transcriptome was conducted among three conditions (control reference, 5 days of storage at 10 °C, and 5 days of storage at 30 °C). The PCC of *Vp* transcriptome between *tdh*+ and *trh*+ strains in the control group was 0.9,9 suggesting that the gene expression pattern of *tdh*+ and *trh*+ strains before storage was similar and could serve as the control reference for following analysis. The PCC of *Vp* transcriptome analysis between the *tdh*+ and the *trh*+ strains after 5-day incubation at 10 °C was 0.94, and the PCC of *Vp* transcriptome between *tdh*+ and *trh*+ strains was 0.89 after 5-day incubation at 30 °C. This indicated that correlation of gene expression pattern between *tdh*+ and *trh*+ strains was reduced at 30 °C compared with 10 °C. Principal component analysis (PCA) of *Vp* transcriptomics data was conducted to examine variances in gene expression among samples. Principal component 1 (PC1) and principal component 2 (PC2) explained 48.69% and 16.89% variances in gene expression of sequenced transcript reads among control/test groups (**Fig. S2b**). The transcriptome of *Vp* at 10 °C were close to counterparts in control condition, and whereas transcriptome of *Vp* at 30 °C were separated from counterparts in control condition. There was slightly more variation in *Vp* transcriptome profiles between *tdh*+ and *trh*+ strains at 10 °C (R^2^ = 0.67) than 30 °C (R^2^ = 0.73) (**Fig. S3**).

Differentially expressed genes (DEGs) at the transcriptomic level were further determined by RNA-seq analysis. Overall, more DEGs were detected in *Vp* transcriptome at 30 °C compared with 10 °C (**Fig. 3**). Specifically, 1,795 and 1,996 DEGs were identified in transcriptomes of *tdh*+ and *trh*+ strains at 30 °C, respectively, and whereas 283 and 984 DEGs were identified in transcriptomes of *tdh*+ and *trh*+ strains at 10 °C, respectively. Among DEGs at 30 °C, 858 and 977 DEGs were significantly upregulated (Log2 fold change ≥ 1 and FDR-corrected *p*-values < 0.05) for *tdh*+ and *trh*+ strains, meanwhile 937 and 1019 DEGs were significantly downregulated (Log2 fold change ≤ −1 and FDR-corrected *p*-values < 0.05) for *tdh*+ and *trh*+ strains, respectively. Among DEGs identified at 10 °C, 139 and 479 DEGs were significantly upregulated (Log2 fold change ≥ 1 and FDR-corrected *p*-values < 0.05) for *tdh*+ and *trh*+ strains, respectively, and 144 and 505 DEGs were significantly downregulated (Log2 fold change ≤ −1 and FDR-corrected *p*-values < 0.05) for *tdh*+ and *trh*+ strains, respectively (17).

**Figure 3.**
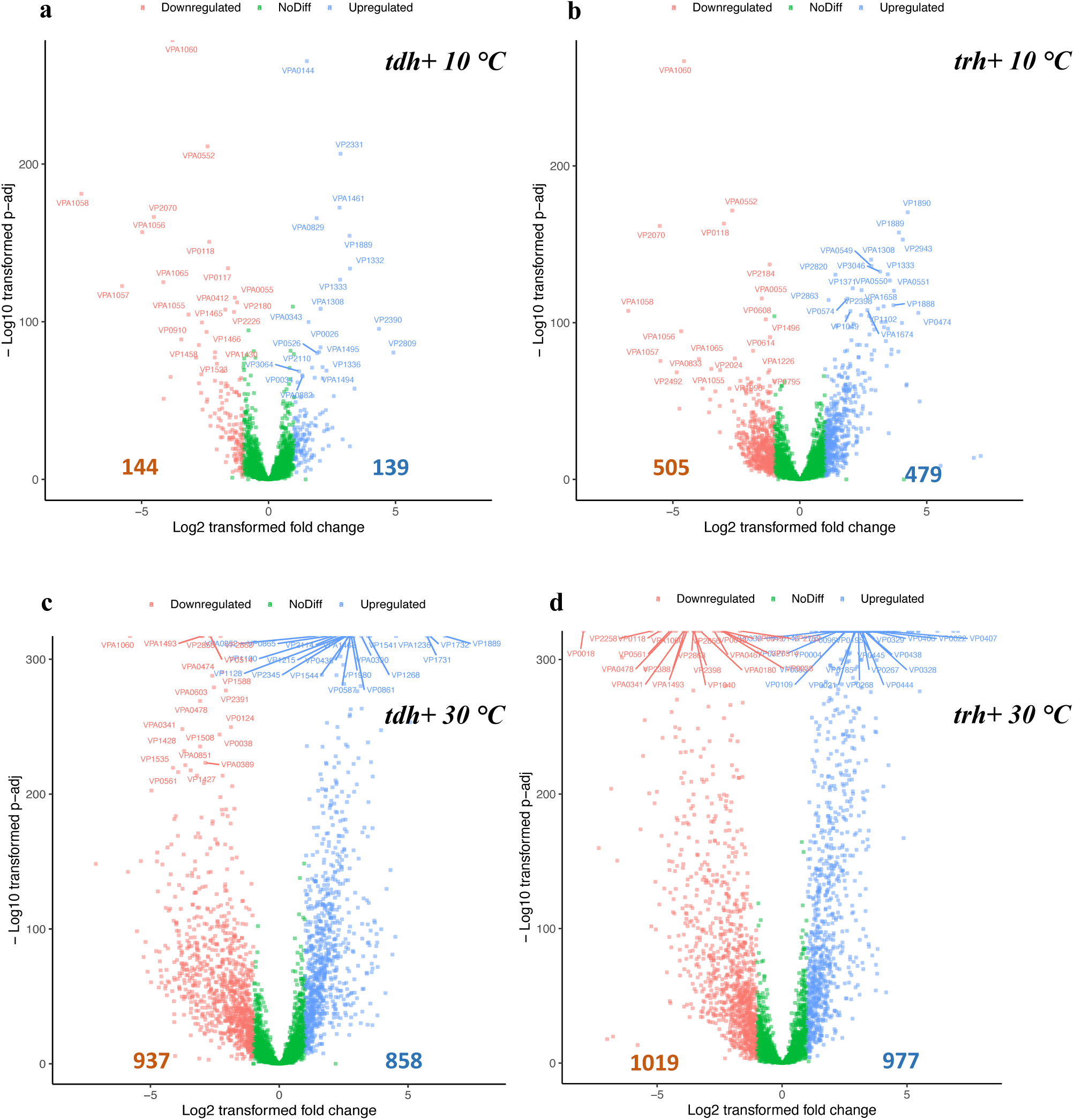
Defferentially expressed genes of *V. parahaemolyticus* with *tdh*+ or *trh*+ genes at 10 (a and b) and 30 °C (c and d) for 5 days. The x-axis represents the log2 of the fold change against the −log10 of the adjusted p-value. Red dots indicate the differentially expressed genes with at least −1.0 change and statistically different from the control (*p* < 0.05). Blue dots indicate the differentially expressed genes with at least +1.0 change and stically different from the control (*p* < 0.05).

To validate the gene expression analyzed by RNA-Seq, seven genes were randomly selected and the gene expression was evaluated using qRT-PCR. Results from qRT-PCR were consistent with that (upregulated or downregulated) of RNA-seq data analysis, suggesting the reliability of results from RNA-seq (**Fig. S4**).

### Biological trace of transition from exponential to stationary phase

For test groups, RNA was extracted from stationary-phase five days after *Vp* incubated at 10 and 30 °C, as it is believed that *Vp* is usually entering into the stationary phase in the environment and food systems (23). When microbial growth shifts from exponential phase to stationary phase, the expression of growth-associated genes becomes slow down and meanwhile persistence-associated genes are increasingly expressed so that bacterial cells can remain metabolically active in stationary phase (24). It is expected that 70S ribosomes are converted into inactive 100S ribosome with loss of ribosomal translation activity via dimerization, which requires ribosome modulation factors (RMF) covering peptidyl transferase domain and the entrance of the peptide exit tunnel (25). For both *tdh*+ and *trh*+ strains, genes annotated to ribonucleoprotein complex and large ribosomal subunit were significantly enriched based on GSEA-GO results (**Figure 7**). Significant activation of cofactor biosynthesis pathway was observed in both *tdh+* and *trh*+ strains based on GSEA-KEGG results (**Figure 9**). These results might suggest the event of ribosome dimerization in *Vp* transitioning from exponential to stationary phase and imply the transition of *Vp* to a non-proliferative metabolic state.

### Cellular response of *Vp* adapting to 10 °C seawater

Although more DEGs were observed in transcriptome of *trh*+ than *tdh*+ strain, less enriched GO terms were enriched through GSEA in *trh*+ strain than *tdh*+ strain. Commonly observed in transcriptome profile of both strains, aromatic amino acid and alpha amino acid groups biosynthesis associated gene clusters were significantly downregulated, indicating *Vp* saved energy budget by avoiding expressing precursors of secondary metabolites **(Fig. 5**). Besides aromatic amino acid biosynthesis significantly enriched functional gene clusters including organic substance transport and cellular amino acid biosynthetic process in *trh* strain were all downregulated, which might suggest the inactive cellular status.

**Figure 4.**
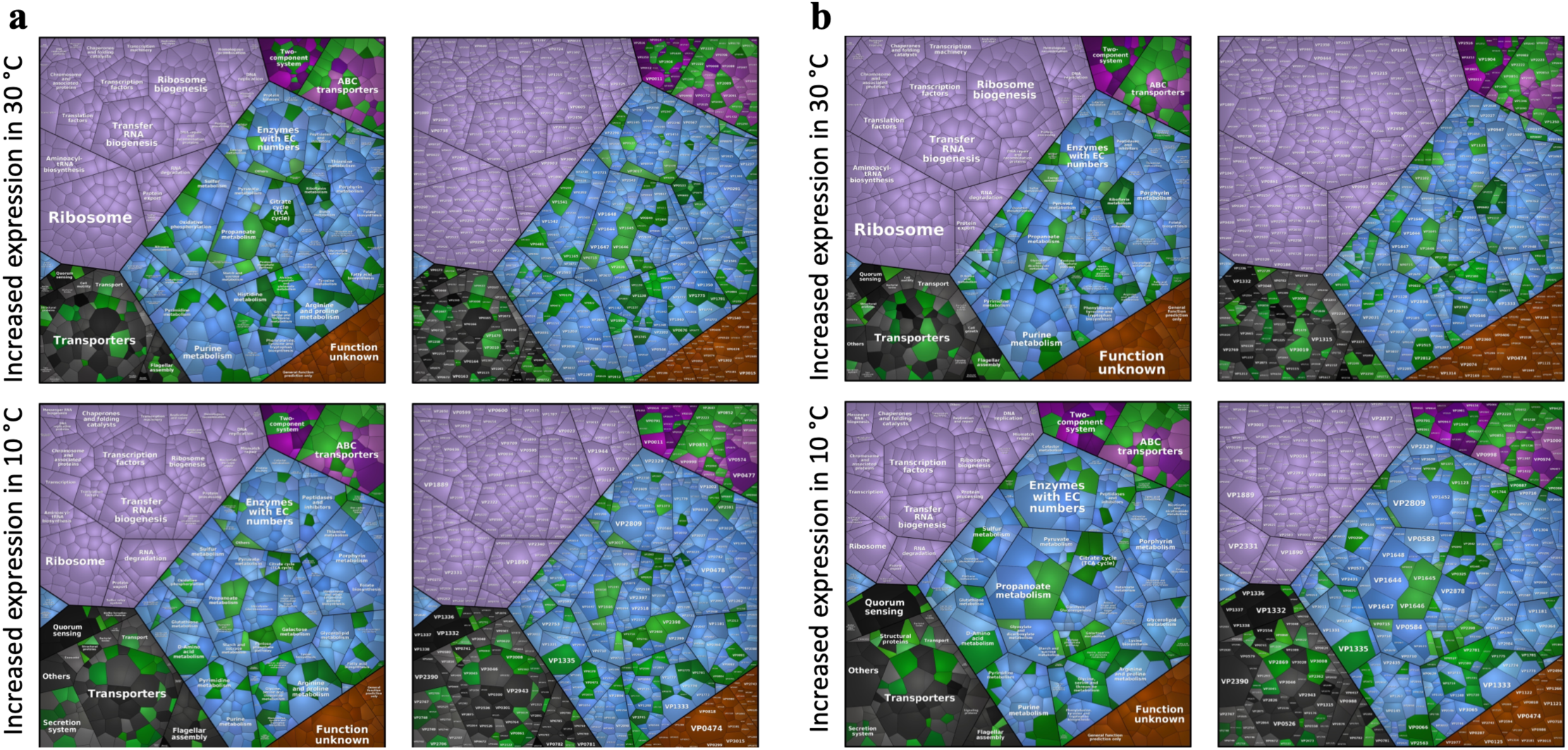
Proteomap illustrating differentially expressed genes of *tdh*+ strain (a) and *trh*+ strain(b) when stored at 10 or 30 °C for 5 days. Genes are clustered by different functional groups.

**Figure 5.**
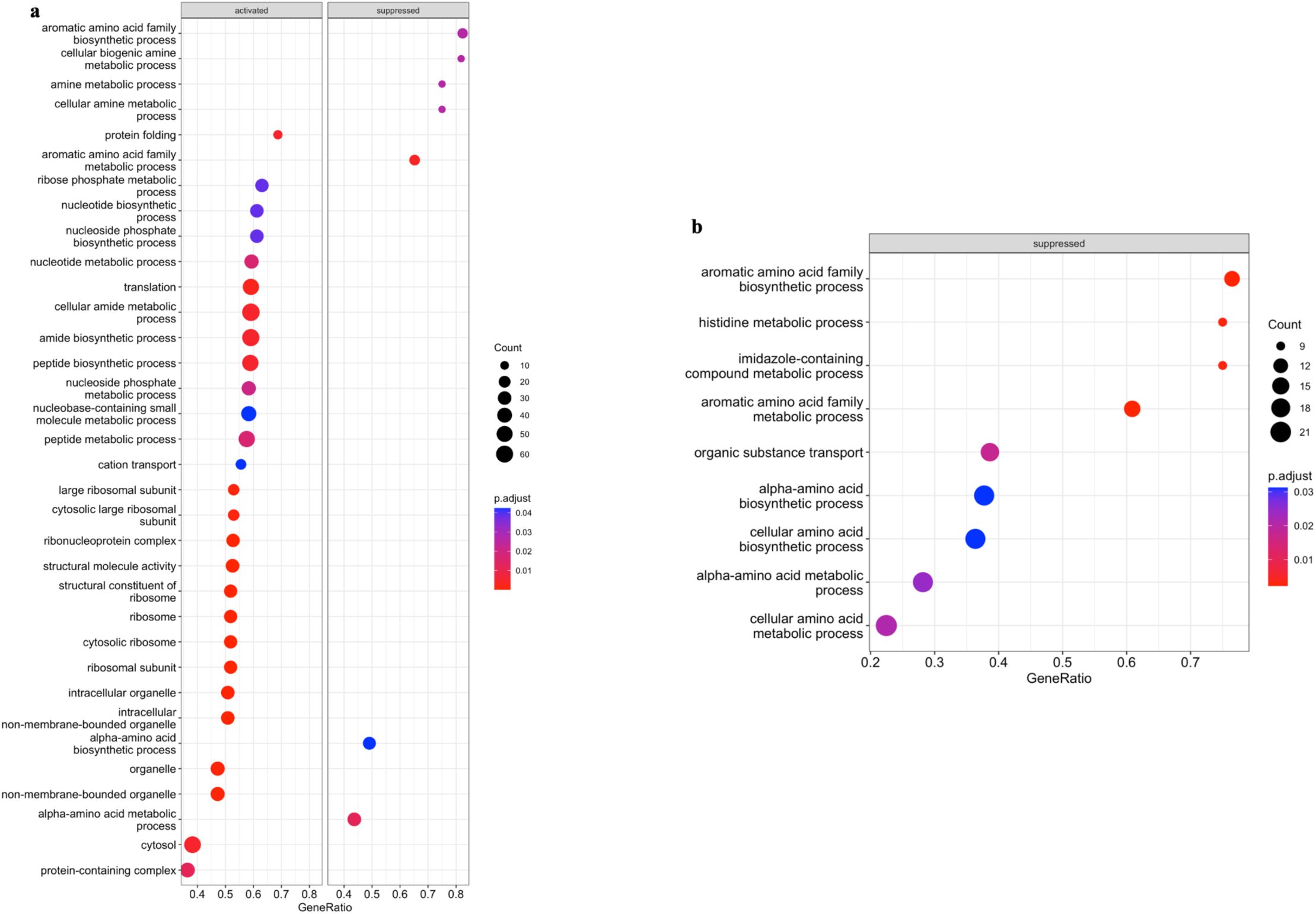
Gene set enrichment analysis against Gene Ontology database (GO) of *tdh+* (a) and *trh+* strains (b) when stored at 10 °C in seawater for 5 days.

Previous studies have indicated that cellular strategies applied by *Vp* to address cold stress include upregulation of cold stress related proteins and increase of membrane fluidity by enhancing fatty acid metabolism (17, 26, 27). Similar results were shown in this study. VP1889 encoding the cold shock protein A (*cspA*) was significantly upregulated for both strains (3.90 and 3.19 Log 2-fold change for *tdh*+ and *trh*+ strains, respectively) at 10 °C (FDR-corrected *p*-values < 0.05). CspA is an RNA chaperone that reduces RNA secondary folding caused by decreasing temperatures. The upregulation of *cspA* indicated that *Vp* counteracted the translational hardness caused by RNA folding at 10 °C by increasing the *cspA* expression (28).

Modification of fatty acid is critical for bacterial survival at low temperatures, as lipid molecules can become more ordered and solidified as temperature decreases (29). T. Xie et al. (17) pointed out that the essential role of co-occurred downregulated pyruvate metabolism and upregulated fatty acid biosynthesis in cold tolerance of *Vp* at 4 °C. Although no direct evidence related to pyruvate metabolism and fatty acid biosynthesis were observed in transcriptome profile of *tdh*+ strain, thiamine metabolism was shown to be significantly upregulated (FDR-corrected *p*-values < 0.05) (**Fig. 7**). Thiamine pyrophosphate (TPP) is the key co-enzyme in fatty acid biosynthesis, which involves in the interconversion of pyruvate to acetal-CoA by pyruvate dehydrogenase, the fundamental precursor in fatty acid biosynthesis (30). The observed upregulated thiamine metabolism pathway might suggest potential upregulated fatty acid biosynthesis in *tdh*+ strain in seawater at 10 °C., pyruvate metabolism was significantly upregulated in *trh*+ strain in seawater at 10 °C (**Fig. 7**). At the proteomic level, J. Tang et al. (31) reported that pyruvate dehydrogenase complex repressor (a regulator negatively impacts the formation of pyruvate dehydrogenase complex, PDHC) was mostly downregulated in *Vp* incubated at 4 °C after 18 hours, and suggested that the resulted enhanced PDHC activity was critical for *Vp* to maintain its viability under cold stresses. Taken together, these results highlighted the active pyruvate metabolism change involved in *Vp* surviving at low temperatures.

### Cellular responses of *Vp* growing at 30 °C

A strong signature of growth was observed in *Vp* incubated at 30 °C compared with 10 °C, including ribosome biogenesis, amino acid metabolism, and purine metabolism (**Fig. 4**). Greater than 50% of DEGs were annotated to biosynthesis processes, including the macromolecule biosynthetic process, cellular biosynthetic process, and organic substance biosynthetic process that were significantly upregulated in *tdh*+ strain (**Fig. 6**). Amino acid biosynthesis pathway and alanine, aspartate and glutamate metabolism pathways were significantly upregulated (**Fig. 7**). Alanine, aspartate and glutamate are critical amino acids serving as precursor for diverse metabolites as essential cellular component for bacterial cell growth (32, 33). In addition, both arginine biosynthesis and arginine metabolism were both significantly upregulated (**Fig. 8**). Similar results were reported in the previous work: L. Li et al. (34) reported that arginine biosynthesis pathway was upregulated in *Vp* incubated in eutrophic outlet water at 30 °C compared with counterparts incubated at 16 °C. Arginine biosynthesis is essential to microbial growth as arginine can be converted putrescine, which serves as an essential regulator for cell growth, differentiation, proliferation, and various physiological processes (35, 36). These upregulated biosynthetic process contributed to the fitness of *tdh*+ strain at optimum growth temperature and maintaining stable populational level at stationary phase.

**Figure 6.**
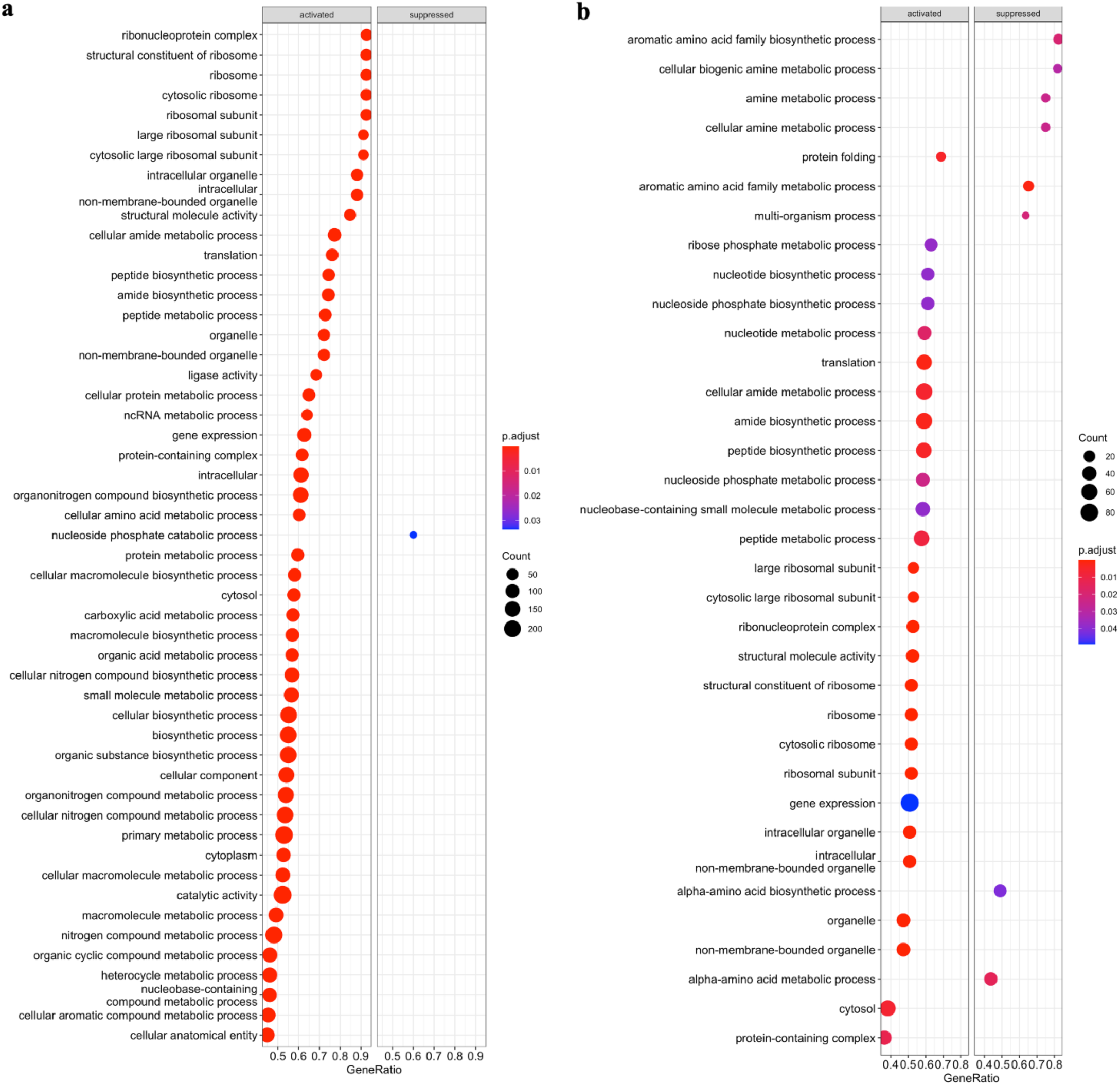
Gene set enrichment analysis against Gene Ontology database (GO) of *tdh+* (a) and *trh+* strains (b) when stored at 30 °C in seawater for 5 days.

**Figure 7.**
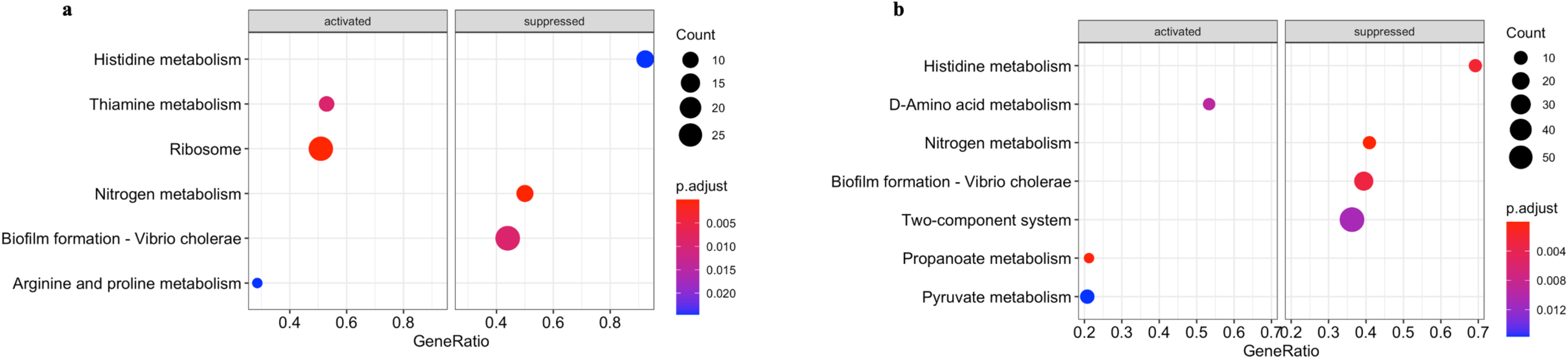
Gene set enrichment analysis against Kyoto Encyclopedia of Genes and Genomes database (KEGG) of *tdh+*(a) and *trh+* strains (b) when stored at 10 °C in seawater for 5 days.

**Figure 8.**
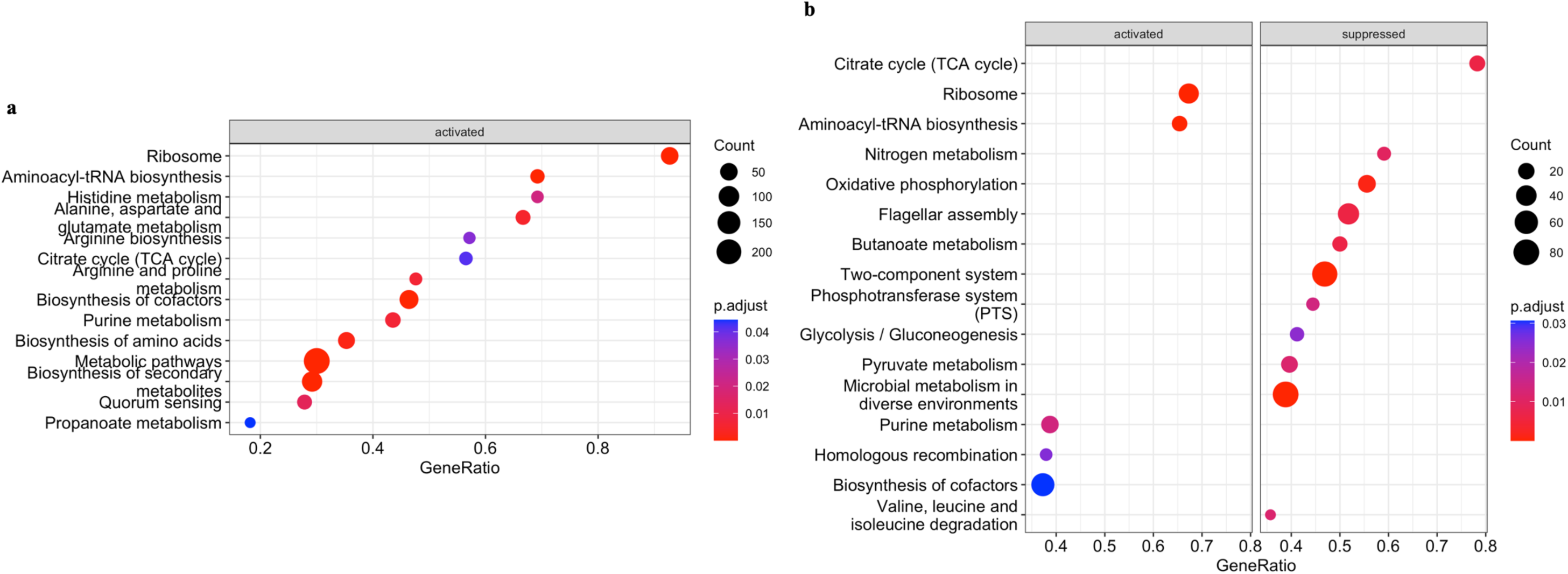
Gene set enrichment analysis against Kyoto Encyclopedia of Genes and Genomesdatabase (KEGG) of *tdh+*(a) and *trh+* strains (b) when stored at 10 °C in seawater for 5 days.

Higher growth rate was observed in *tdh*+ strain in comparison to *trh+* strain at the phenotypic level. Results at transcriptomic levels provided additional biological insights. More significantly downregulated functional gene clusters and metabolism pathways were detected in *trh+* strain than *tdh*+ strain based on GSEA results (**Figs. 6 and 9**). Functional gene clusters associated with ribosome, ribosome biosynthesis, and transfer RNA biosynthesis coupled with pathway enrichment in central energy metabolism reflect energy use for cell growth and proliferation of *tdh*+ strain at 30 °C. Central energy metabolisms include glycolysis/gluconeogenesis, pyruvate metabolism, and TCA cycle were significantly downregulated in the *trh+* strain after 5 days of incubation at 30 °C, suggesting that the energy generation was weakened in *trh+* strain. Moreover, oxidative phosphorylation pathway was significantly downregulated in *trh+* strain as well, indicating the process of intracellular ATP synthesis was inhibited. In addition to downregulation of energy metabolism pathways, significantly downregulated usage of valine, leucine, and isoleucine pathway was detected in *trh+* strain (**Fig. 8**). Valine, leucine, and isoleucine are the core branched chain amino acids essential for bacterial growth (37). Such decreased degradation of branched chain amino acid might inhibit interconversion to metabolites essential for growth and co-factors. This observed *trh+* strain-only inferiority due to turned off central energy metabolism might explain its lower fitness in comparison to *tdh*+ strain in seawater at 30 °C.

### Expression of virulence genes at different temperatures

Microbial pathogenesis can be significantly affected by environmental temperature (38). Virulence and pathogenesis of *Vp* was commonly reported based on studies using live animal models (39–41). However, limited information about its virulence in natural seawater environment has been reported at this moment. Based on the GSEA-KEGG results, the biofilm formation pathway was significantly downregulated at 10 °C; this has been observed in both *tdh*+ and *trh*+ strains. N. Han et al. (42) reported that the biofilm formation of *Vp* on food and food surfaces increased as the environmental temperatures increased. Both results highlight the importance of temperature during both pre-harvest production and handling and post-harvest processing and storage. Moreover, histidine metabolism was significantly downregulated in both strain *Vp* at 10 °C, indicating potential decreases in histamine formation (**Fig. 7**). Histamine is a major allergen in aquaculture products (43). Less active cellular status of *Vp* in seawater during winter season could lead to less metabolites potentially serving as causative allergic agent to human. To validate this observation, metabolomic profiling of *Vp* in seawater at different temperature warrant additional future studies.

S. Urmersbach et al. (26) reported that expression of major virulence-associated genes of *Vp* RIMD2210633 such as *tdh*, *tox*R, *tox*S remained unaffected by cold and heat shock in alkaline peptone water (4 and 42 °C, respectively). In this study, our results showed that virulence-associated genes, including *toxR*, *toxS*, and T3SS1 effectors *vopQ*, *vopR*, *vopS*, and VPA0450, showed less than 1.0 Log 2-fold change. No significant gene expression of VPA1509 was found in *tdh*+ and *trh*+ strain at 10 °C over 10-day storage. The less active expression of major haemolysin (VPA1509) in *Vp* in cold might be due to temperature dependent conformational change in *tdh* and *trh* encoded hemolysin protein, which led to increasing energy cost of transcription (44). The expression level of haemolysin encoded genes was significantly downregulated 30 °C (with −1.99 and -2.14 Log 2-fold change for *tdh*+ and *trh*+, respectively). Repressed expression of virulence-associated genes was expected when bacteria need to actively regulate genes coding for enzymes essential for growth in the optimal environment (45).

Moreover, VP1890 (*vacB*) encoding a putative virulence-associated protein was significantly upregulated for *tdh*+ and *trh*+ strains at both 10 and 30 °C (with 4.25 and 2.13 Log 2-fold changes for *tdh*+ and *trh*+ strains at 10 °C, respectively; 4.27 and 3.42 Log 2-fold change for *tdh*+ and *trh*+ strains at 30 °C, respectively, FDR-corrected *p*-values < 0.05). The product of *vacB* was reported to be an exoribonuclease RNase contributing to the virulence of *Shigella* and enteroinvasive *Escherichia coli* (46). The strong expression of *vacB* in *Vp* was consistent with previous studies. L. Meng et al. (47) reported more than 5 Log 2-fold upregulation of *vacB* in viable but non-culturable state *Vp* induced at 4 °C over 40 days. S. Urmersbach et al. (26) also reported *vacB* showed the highest upregulation (7.01 Log 2-old change) in *Vp* when being incubated in APW at 15 °C for 30 min. All of combined information confirmed high expression levels of *vacB* at temperatures ranging from 4 to 15 °C. In this study, the expression of *vacB* was also upregulated at 30 °C. As discussed by Liao et al., the detection of *Vp* at lower temperatures can be challenging, newer candidate genes that continuously expressed at higher levels at various tempeatures are needed. Through this study, results indicated that *vacB* could be served as a potential biomarker to identify *Vp* in natural coastal environment across seasons. Through BLAST, the *vacB* nucleotide sequence shows high specificity in *Vp*.

## CONCLUSION

This study investigated the fitness and cellular response at the transcriptomic level of two *Vp* strains (*tdh*+ and *trh*+) in seawater at different temperatures corresponding to oyster harvesting seasons (10 and 30 °C). When *Vp* was incubated in seawater at 10 °C, persistence of both *tdh*+ and *trh*+ strains was observed over 10 days and higher die-off rate was observed on *trh*+ than *tdh*+ strain based on predicted model. When *Vp* was incubated in seawater at 30 °C, growth of *tdh*+ strain was better than *trh*+ strain and higher growth rate was observed on on *tdh*+ than *trh*+ strain based on predicted model. More DGEs were detected at 30 °C than 10 °C, indicating that cellular responses of *Vp* at the transcriptomic level were more complex during summer months. Expression of cold shock associated genes in *Vp* in seawater at 10 °C were upregulated. No remarkable gene expression of VPA1509 (*tdh*1) was observed in *tdh*+ and *trh*+ strains at 10 °C over 10-day storage, but the gene expression was significantly downregulated 30 °C, which highlighted cost-effective energy allocation strategy of *Vp* during growth and persistence. The *vacB* gene encoding a putative virulence-associated protein (VP1890) presented significantly upregulated expression in *tdh*+ and *trh*+ strains at both 10 and 30 °C. In addition, *tdh*+ and *trh*+ strains in seawater at 10 °C showed downregulated biofilm formation pathway and histidine metabolism. Based on biological insightfulness from *Vp* transcriptome profile, pre-harvesting temperatures play impacts on the cellular response and virulence of *Vp* in seawaters. The valuable information provided in this study reveal that it is critical to understand behaviors of *Vp* to better assist with *Vp* risk assessment and management in oyster harvesting season.

## MATERIALS and METHODS

### Culture preparation and incubation conditions

Frozen culture of *Vp* strains ATCC 43996 (*tdh*+) and ATCC 17802 (*trh*+) purchased from American Type Culture Collection (ATCC) were streaked and activated on Tryptic Soy Agar (TSA) supplemented with 3% NaCl. Cultures were incubated at 37 °C for overnight. After that, a single colony was picked from each TSA plate and transferred into 10 ml of Tryptic Soy Broth (TSB) supplemented with 3% NaCl for additional 24 hours of incubation at 37 °C. After incubation, a loopful of fresh liquid culture was transferred into another 10 ml of fresh TSB supplemented with 3% NaCl. The inoculated TSB was incubated at 37 °C for overnight and washed on the next day by centrifugation (Eppendorf, Hauppauge, NY, USA) at 3,000 × g for 10 min. Washed cultures were resuspended with ca. 2 ml of phosphate-buffered saline (PBS; pH 7.4). The optical density at 600 nm (OD600) of each washed culture was adjusted to 1.6 ± 0.1 by using a spectrophotometer (Thermo Scientific, Piscataway, NJ, USA). This washed and adjusted culture has approximate 7.0 Log CFU/ml of cells based on plate count results. To inoculate the seawater, 1 ml of washed culture was added into 9 ml of autoclaved natural seawater. Seawater was collected from the from the Auburn University Marine Extension and Research Center located in Dauphin Island, AL. Inoculated seawater samples were first kept at ambient temperature for 2 hours to enable the *Vp* cultures to adapt to the new environment. After 2 hours, inoculated seawater samples were stored at 10 °C for 10 days or 30 °C for 5 days.

### Enumeration of *Vp* in inoculated seawater samples by using the plate count method

During storage, sub samples (1 ml each) were taken and plated every 24 h for the 10 °C storage condition and were plated every 2 h for the first 12 h then at hours 24, 48, 72, and 120 for the 30 °C storage condition. Three biological replicates were conducted. Every 1 ml of seawater sample were diluted in serial 10-fold dilutions, and plated onto Thiosulfate-citrate-bile salts-sucrose (TCBS) plates (BD, Sparks, MD, USA). Plates were incubated at 37 °C for 18 h before enumeration. Plates were then placed back to the incubator and the colony counts were confirmed after another 24 hour of incubation. *Vp* concentrations were expressed in common logarithm transformation format with the unit of CFU/ml.

### Statistical analysis and predictive models for describing *Vp* behaviors in seawater

The populations of *Vp* present in seawater were enumerated at different time intervals at 10 and 30 °C. One-way Analysis of Variance (ANOVA) followed by the Tukey test was applied to compare the difference in *Vp* concentrations as predicted by primary predictive models. Primary predictive models were established for two *Vp* strains at two storage temperatures with the OriginPro 2023 software (OriginLab Corporation, Northampton, MA, USA). *P* < 0.05 was considered statistically significant. The Baranyi model (see equation 1) was chosen to fit the *Vp* population data and the calculations were performed using the DMfit tool available at the Combase website, https://browser.combase.cc/, (48). The equation of Baranyi model is as follows:

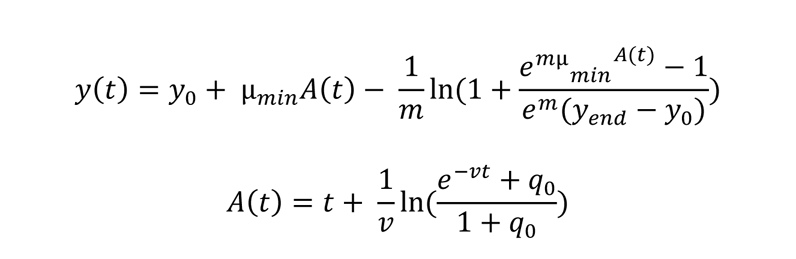

where *y* is the natural logarithm of the bacteria concentration at any given time (ln CFU per milliliter), *y*_o_and *y*_end_are the initial value and the end value of y, *A*(*t*) is the equation governing the duration of the period preceding the Log linear inactivation phase, *t* is time (day), *m* determines the smoothness of the transition from the exponential inactivation phase to the survival tail, μ_min_ is the minimum value of the inactivation rate or the maximum value of the growth rate, *v* is the rate at which the bacteria lose the ability to survive during the shoulder, and *q*_o_ is the initial physiological state of bacterial cells.

### RNA extraction and sequencing

RNA samples extracted from the *Vp* 2 hours after inoculation was labeled as the control, and RNA samples extracted from *Vp* strain at the end of 5 days of storage at 10 or 30 °C was labeled as test group. To extract the RNA, 1 ml of *Vp* culture was taken and centrifuged at 3,000 x g for 10 min. The supernatant was removed and the cell pellet was re-suspended in 1 ml of PBS. The total bacterial RNA was extracted by using the Qiagen RNeasy mini kit (Qiagen, Valencia, CA) following the manufacturer’s instruction. The quality of the extracted RNA was measured with the Agilent 2100 electrophoresis bioanalyzer (Agilent, Santa Clara, CA) to ensure that the RNA integrity numbers (RIN) of all RNA samples were greater than 7.0. Once the RNA quality was confirmed, cDNA library was prepared by using the QuantiTect reverse transcription kit (Qiagen, Valencia, CA). The cDNA library was then sent to the Genomic Services Laboratory at HudsonAlpha Genome Sequencing Center (Huntsville, AL) and sequenced on the Illumina HiSeq 2500 platform to generate 50 bp pair-end reads.

### Transcriptomics analysis

A schematic illustration of transcriptomics analysis pipeline is shown in **Fig. 1**. The quality of raw reads was checked by FASTQC. Given the small size of bacterial genome (approximately 5.1 MB for *Vp*) and since there was no potential mRNA splicing issues, a fast and bias-aware analytical pipeline, Salmon, was used to achieve quantification of transcript expression mapping (49, 50). Salmon was used to align the reads against the *Vp* RIMD 2210633 protein coding sequence (CDS) region of each gene (reference transcriptome) [GenBank accession number GCA_000196095.1]) with parameters -gcBias. Read counts were imported into R, filtered, and converted to mRNA transcripts by using the tximport package (51). Deseq2 was used to achieve data normalization and identify differentially expressed genes (52). Genes with an FDR-corrected p-values < 0.05 were considered significant. Threshold of Log2 fold change ≥ 1 was considered upregulated and threshold of Log2 fold change ≤ 1 was considered downregulated (17). Reference transcriptome annotation against Gene Ontology database was conducted by using eggnog-mapper (53). Gene set enrichment analysis (GSEA) was performed for genes that were upregulated and downregulated in both storage temperatures against the Gene Ontology (ont = ALL) and the Kyoto Encyclopedia of Genes and Genomes (organism = vpa) databases by using the R package clusterProfiler (54). Differentially expressed genes were visualized by using proteomaps (55). The reference gene list of *Vp* used in proteomaps was constructed based on the JSON file in the KEGG orthology database by Python (treemap template ID: *Vibrio parahaemolyticus* RIMD 2210633 V7).

### Quantitative real-time PCR (qRT-PCR) validation

To validate the transcriptomics results, genes of each *Vp* strain showing the same up-or down-regulation patterns at both 10 and 30 °C were selected for validation studies. The gene *pvuA* was used as the housekeeping reference gene as its expression level has shown to be stable through a wide range of temperatures (56). The expression level of the select gene was normalized to reference gene and calculated by using the −ΔΔCt method (57). The list of primers used in this study is provided in supplementary **Table 1**. qRT-PCR was conducted on the QuanStudioTM Real-Time PCR system (Applied Biosystems, Foster City, CA). The total volume of each reaction was set to 25 μl, consisting of 2 μl cDNA aliquot (concentration 1ul/ml), 1 μl forward primer (1 μm/μl), 1ul reverse primer (1 μm/μl), 12.5 μl 2 × SYBR™ Green PCR Master Mix (Life Technologies, Carlsbad, CA, United States), and 8.5 μl nuclease water. Program of thermocycler was set to start with an initial denaturing period at 95 °C for 10 min and then 40 cycles of 95 °C for 15 s, 52 °C for 20 s and 72 °C for 25 s. The specificity of the PCR product was checked by analyzing the melt curve.

## Data availability

The raw and processed RNA-Seq data can be found in the GEO database under accession number PRJNA949728 and PRJNA949727.

## SUPPLEMENTAL MATERIAL

Supplemental material is available online only.

## ACKNOWLEDGMENTS

This work was supported by funding from USDA HATCH grant S1077.

L.W. and C.L. designed the study. C.L. and Z.L. performed the experiments. C.L. analyzed the *Vp* fitness data. Z.L. analyzed the RNA-Seq data. Z.L. initiated the manuscript draft.

C.L. and L.W. reviewed and revised the manuscript.

We declare no competing interests.

---

**Figure S1.**
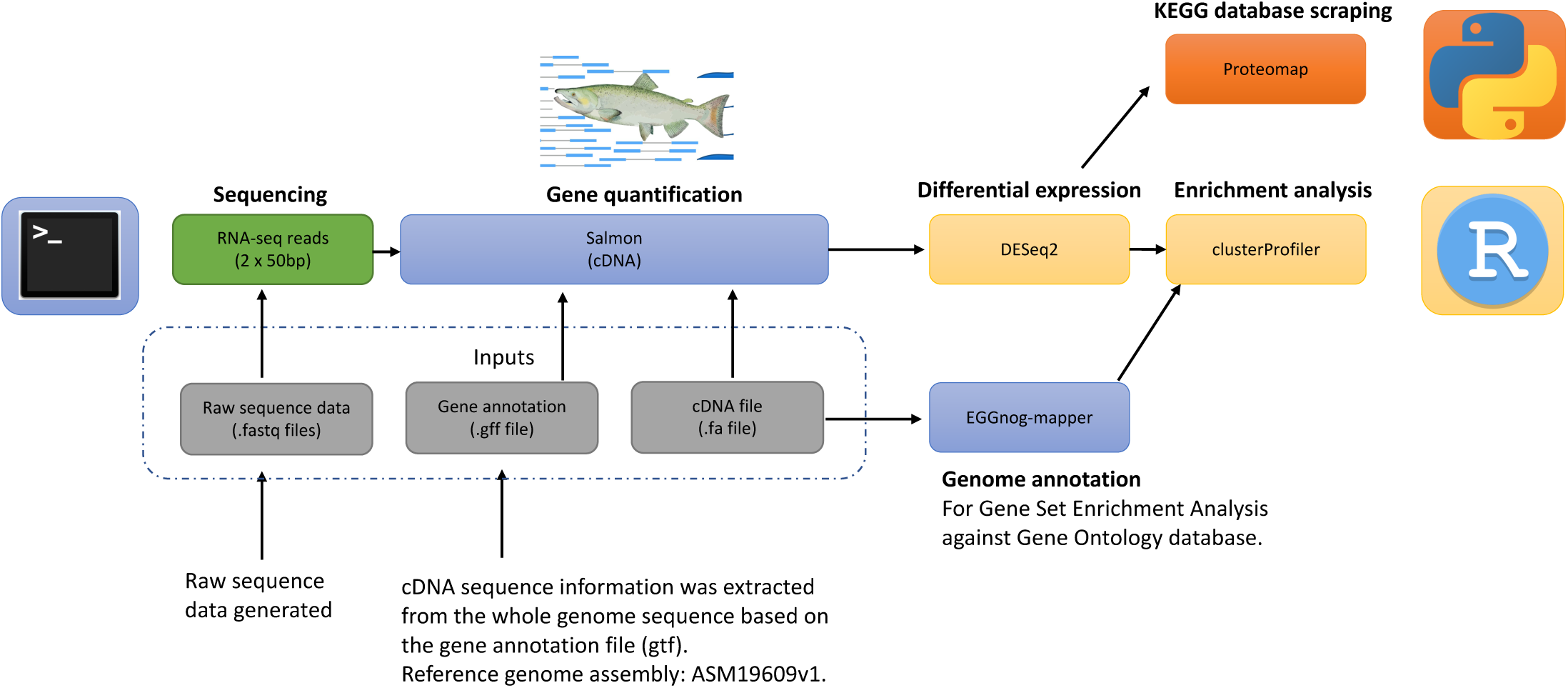
A schematic illustration of the analytical pipeline for the transcriptomic data. Blue color indicates steps conducted in linux system. Yellow color indicates steps conducted in R. Orange color indicates steps ucted in Python.

**Figure S2.**
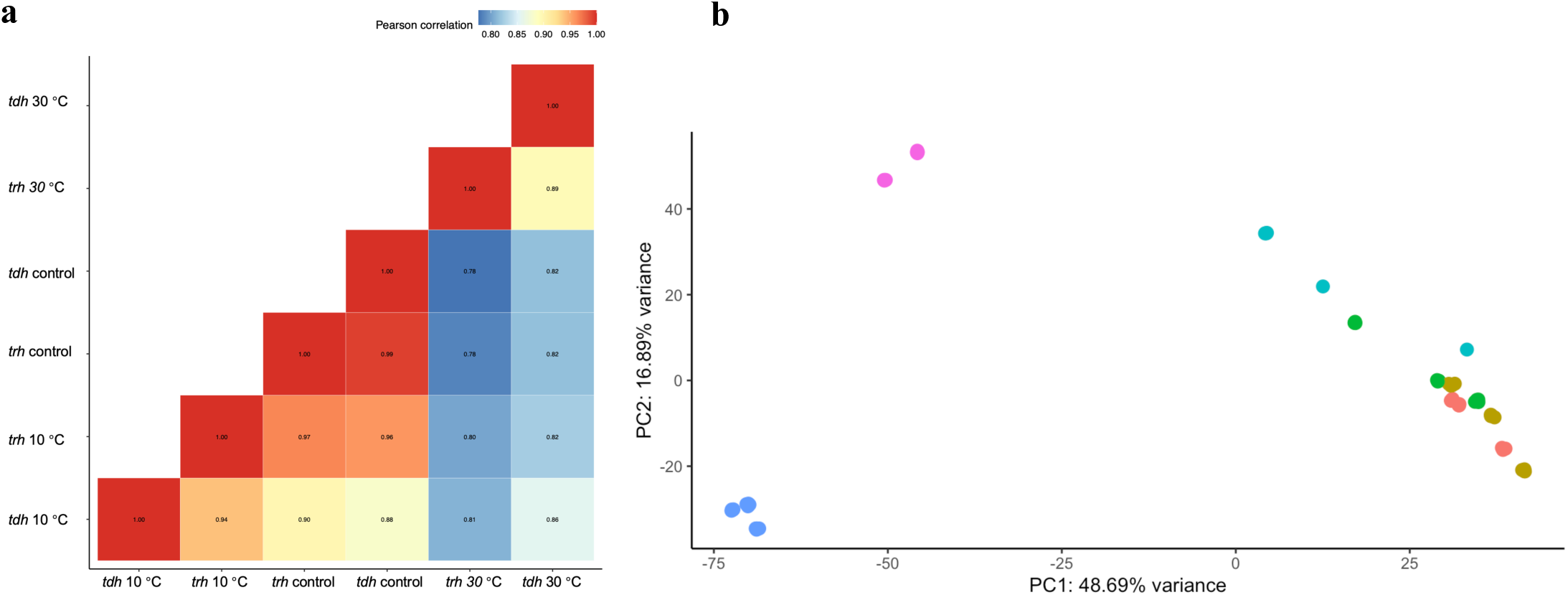
Pearson correlation (a) of transcriptomic profiles *V. parahaemolyticus* in the control group (2 hours inoculation) and tested groups (10 and 30 °C); Principal component analysis (b) of transcriptomic profiles *V. arahaemolyticus* in the control group (2 hours after seawater inoculation) and tested groups (10 and 30 °C). 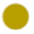t*dh*+ and 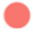*trh*+ in the control group, 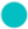*tdh*+ and 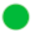*trh*+ in 10 °C group, and 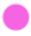*tdh+* and 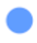*trh+* in 30 °C group.

**Figure S3.**
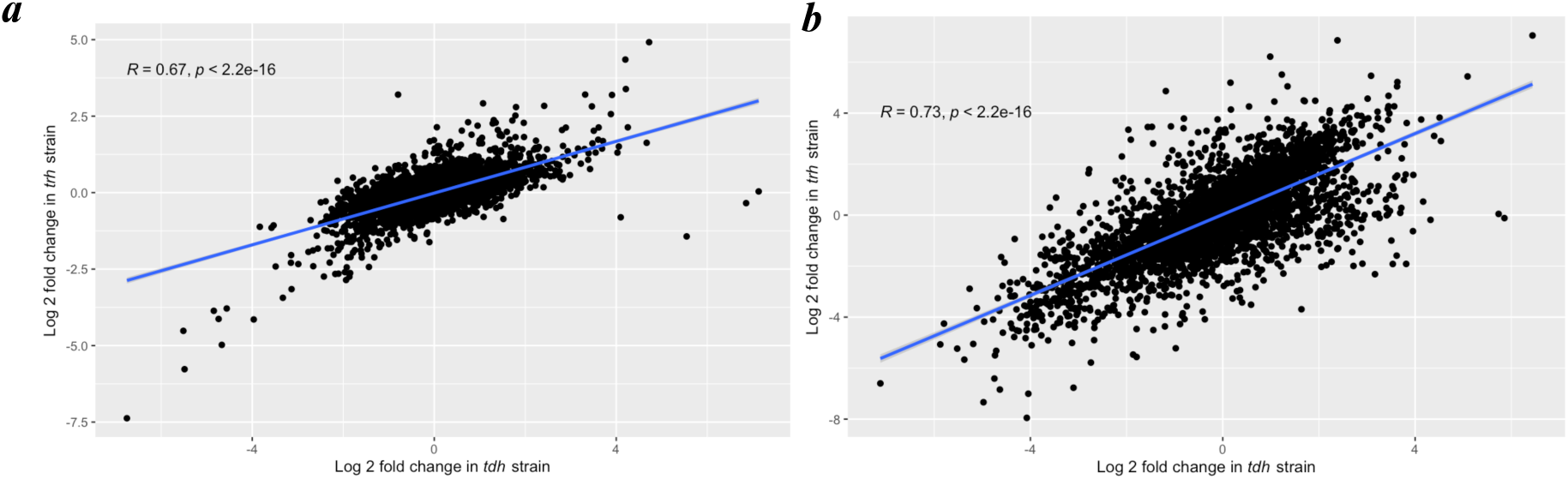
Scatter plots of the comparison of gene expression levels between *tdh*+ and *trh*+ strains when stored at °C (a) and 30 °C (b) for 5 days.

**Figure S4.**
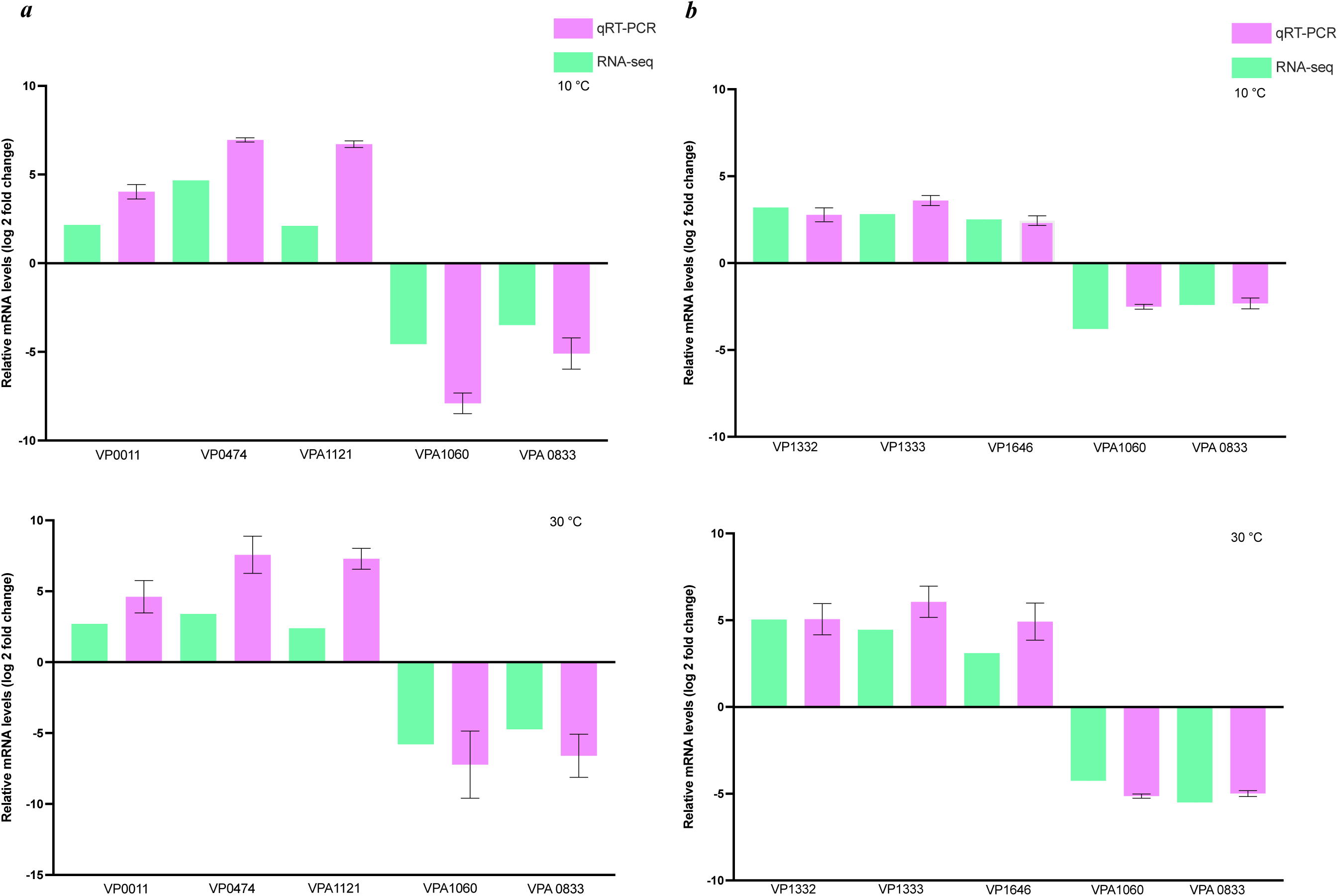
Validation of the select differentially expressed genes identified from RNA-seq results by qRT-PCR for *tdh*+ strain (a) and *trh*+ strain (b).

